# Floral nectar microbiota of *Persea americana* inhibits pathogens and improves plant fitness

**DOI:** 10.1101/2025.03.27.645287

**Authors:** Claudia Marina López-García, Indira Aranza Rodríguez-Gómez, Yareli Pérez-Bautista, Luis Alberto Villanueva-Espino, Mariana Molina Torres, Violeta Patiño-Conde, Luis Enrique Ruiz-Guizar, Mariel García-Meléndez, Orlando Hernández-Cristóbal, Jesús Llanderal-Mendoza, Mauricio Quesada-Avendaño, Frédérique Reverchon, Ken Oyama, Alfonso Méndez-Bravo

## Abstract

The floral nectar-living microorganisms contribute to flower protection and influence mutualistic relationships between plants and pollinators. Avocado floral nectar is poorly attractive to its introduced pollinator *Apis mellifera*. More than 99% of the flowers produced by avocado trees are not able to set fruits, falling to the ground; the interaction between floral microbiota with plant roots is an emerging ecological topic to study.
Here, we examined the richness and abundance of bacteria and fungi from avocado nectar and screened the culturable fraction for their antagonistic activity against the avocado pathogens *Phytophthora cinnamomi* and *Colletotrichum gloeosporioides*, and against the most devastating honeybee pathogens *Ascosphaera apis* and *Paenibacillus larvae*. Experimentally, we also analyzed the effects of the microbial isolates on plant growth and the activation of the jasmonic acid (JA) defense-responses in the model plant *Arabidopsis thaliana*.
*Pseudomonas*, *Acinetobacter*, *Protomyces* and *Vishniacozyma* were the dominant genera. From 43 isolates evaluated, 20 bacteria, and three yeasts showed differential inhibitory activity against the plant and honeybee pathogens, promoted the growth of *A. thaliana* seedlings and induced JA signaling.
Collectively, our findings highlight the selectivity of avocado nectar over its microbiota, which could directly impact the plant fitness and contribute to its pollinatorś health.

**Highlights:** Floral nectar microbes from avocado inhibit pathogens and improve plant fitness.

## 1. Introduction

Pollination in angiosperms is a crucial event for the conservation of terrestrial ecosystems that involves a multipartite ecological interaction between flowers, pollinators and their associated microorganisms (Vannette, 2020). Nectar-consuming insects shape the composition of microbial communities in flowers, which could result neutral, beneficial or pathogenic for plants (Mueller et al. 2022). In turn, the floral microbiota alters nectar chemistry and influences pollinator and plant health (Cullen et al. 2021; Martin et al. 2022). Nectar, a substance present in flowers, mainly contains sugars, amino acids and secondary metabolites and can be considered a niche where only microorganisms tolerant to osmotic stress and low nitrogen availability can develop (Warren et al. 2020). Nectar microbial communities tend to be species-poor and dominated by a few bacteria and yeast taxa (Mueller et al. 2022). Microbial isolates present in nectar can range from plant, pollinator and even human pathogens, to potential commensal and attractors of pollinators and natural enemies of pest insects and phytopathogens (Sasu et al. 2010; Adler et al. 2021; Álvarez-Pérez et al. 2024), positioning nectar as a potential source of beneficial microorganisms for the plant-pollinator interactions and inhibition of plant pathogens. However, the search for pathogen antagonists and plant growth-promoting or defense-inducing microorganisms from nectar has been scarcely studied.

*Persea americana* Mill. (avocado) (Lauraceae), an ancient angiosperm native from Mexico and Central America has gained great relevance worldwide due to its nutritional value (Rendón-Anaya et al. 2019; Bhore et al. 2021). The agricultural management of this crop is highly input-intensive due to the incidence of fungal diseases and a low pollination rate, among others (Pérez-Jiménez 2008; Twizeyimana et al. 2013). Among the most impactful diseases is anthracnose, caused by the opportunistic phytopathogen *Colletotrichum gloeosporioides* (Xoca-Orozco et al. 2017). Over the past five years, major avocado producing countries have reported phytosanitary alerts because the *C. gloeosporioides* incidence in the orchards (Ayvar-Serna et al. 2021; Bustamante et al. 2022). Although anthracnose is a disease that manifests in fruits, infection usually occurs within the flowers, which hosts bacteria and fungi that inhibit the growth of several *Colletotrichum* spp. pathogens in blueberries (Rering et al. 2023). Therefore, the potential of nectar microorganisms to act as biocontrol agents of *C. gloeosporioides* in avocado should be investigated.

Low pollination rates occur in avocado for several reasons. The small, greenish avocado flowers open twice; first in the pistillate female stage in the morning and in the staminate male stage in the following afternoon, or the opposite flowering rhythm, the female stage in the afternoon and the male stage in the following morning (Bender 2002; Can-Alonzo et al. 2005), thus relying on pollinators for the transfer of pollen from male-stage flowers to stigmas of female-stage flowers (Dymond et al. 2021).

Furthermore, these flowers produce a peculiar nectar with a high sucrose concentration (79-80%), a small fraction of fructose (1%), traces of K+, stachyose and perseitol (Liu et al. 1995). Floral phenology in avocado (dichogamous flowering pattern, low pollen release and widespread pollinator declines) (Dymond et al. 2021) combined with the highly specific composition of the avocado nectar, means that less than 1% of the flowers reach the fruit set (Bender 2002) and the rest falls and decomposes, probably incorporating their microbiota into the soil. Therefore, the potential of avocado nectar microorganisms to interact with the microbial soil communities or colonize the rhizosphere should be elucidated to harness some of the nectar microbiota agroecological benefits. A systematic prospection of nectar microbial communities and their ability to improve the plant fitness is thus crucially needed.

One of the strategies implemented to remedy the low pollination rate in avocado has been the introduction of *Apis mellifera*, impacting positively the production in commercial orchards (Peña and Carabalí 2018). Unfortunately, the management of this generalist pollinator has caused the emergence of pathogens such as the bacterium *Paenibacillus larvae* (Fünfhaus et al. 2018) and the fungus *Ascosphaera apis* (Chaimanee et al. 2017), leading to bee colony losses (Genersch 2010; Goulson et al. 2015). In this context, it is important to determine whether *P. larvae* or *A. apis* are part of the avocado nectar microbiota or whether nectar microorganisms could antagonize the growth of these bee pathogens.

The high selectivity of floral nectar as a microenvironment and resulting competitiveness of its microbiota (Warren et al. 2020; Mueller et al. 2022), combined with the close contact between nectar-living microorganisms and plant tissues (Sasu et al. 2010), led us to hypothesize that nectar microbiota 1) performs probiotic functions in their plant hosts and pollinators as antagonists of pathogens, 2) competes with soil microorganisms when flowers are shed, 3) possesses phytostimulating properties influencing endogenous plant signaling, and that 4) avocado nectar, with its particular chemical composition, recruits a different microbial community than that described in other plant models. Here, we first used a culture-dependent approach to characterize the antagonistic activity of avocado nectar microbiota against the avocado pathogens *C. gloeosporioides* and *Phytophthora cinnamomi* and the honey bee pathogens *A. apis* and *P. larvae.* Moreover, we tested their ability to promote plant development and defense in the model plant *Arabidopsis thaliana*. Secondly, we used metabarcoding sequencing to explore the diversity and composition of the avocado nectar microbial community. Collectively, this information could help us design sustainable strategies to improve avocado production, whilst contributing to the understanding of multipartite interactions between pollinators, plants, and their associated microbiota.

## 2. Materials and methods

### 2.1. Study site and sampling

The study site was located at Acuitzio del Canje, state of Michoacán, Mexico (19°30’ N; 101°20’ O) at 2,080 m.a.s.l., in the main avocado producing area worldwide (FAOSTAT 2016). The neighboring vegetation corresponds to oak forests. Rainfall ranges between 800 and 1,300 mm year^-1^, with an average annual temperature of 24 °C, a maximum temperature of 32 °C and a minimum of 10 °C. Avocado nectar samples were collected with sterile 5 µL capillaries by random sampling of at least 10 female-phase flowers from five different avocado trees without symptoms of any disease in March 2019. Samples were transported for microbiological or molecular processing in the laboratory.

### 2.2. Isolation of the nectar culturable microbial fraction

Nectar samples were immediately mixed with 100 µL of 0.9% sodium chloride upon arrival at the laboratory. Suspensions were spread on the surface of Petri dishes with solid R2A medium supplemented with 20% sucrose and incubated at 28 °C. Isolated colonies were re-streaked on Luria-Bertani (LB) solid medium and potato dextrose agar (PDA) supplemented with 100 mg L^-1^ of ampicillin until pure cultures were obtained. The 97 resulting microbial isolates were grouped into 43 morphotypes, according to morphological criteria and Gram-staining as described in Guevara-Avendaño et al. (2019). All isolates were preserved at -20 °C in LB or potato dextrose broth supplemented with 20% glycerol.

### 2.3. Antagonism assays against avocado pathogens

Avocado pathogens were the fruit fungal pathogen *C. gloeosporioides* and the soil-borne oomycete *P. cinnamomi*. To implement the dual antagonism assays, 10 µL of a standardized suspension (0.5 of absorbance at OD600) from each nectar microbial isolate were inoculated at 2.5 cm from a mycelium disk (5 mm-diameter) placed in the center of PDA plates. Three isolates were evaluated per plate, as control, 10 µL of LB medium were inoculated, following the design described in Guevara-Avendaño et al. (2018). To evaluate the antagonistic activity of strains by volatile emission, the two- sealed-base-plates method described in Guevara-Avendaño et al. (2019) was employed. Briefly, the strains were pre-grown for 24h as described above, lids were replaced by another base plate containing a disc of 5 mm diameter of fungal or oomycete mycelium on PDA. The plates were sealed one each other with Parafilm® and incubated at 28 °C. Three assays were set up only with growth media as controls. After 5 days of incubation, the fungal growth inhibition percentage was determined by measuring the mycelial diameter, with the formula described in Guevara-Avendaño et al. (2018). Each combination of isolates was tested in triplicate. To assess the damages induced in the hyphae, 10 representative mycelium samples of each tested fungus/oomycete grown under control conditions and in the presence of the two isolates with the highest antifungal activity were collected, processed and analyzed in JEOL- IT300LV scanning electron microscope and software JSM-IT300 v. 1.020 Scanning Electron Microscopy (SEM) according to Méndez-Bravo et al. (2018).

### 2.4. Molecular identification of nectar isolates

DNA was extracted from the nectar isolates which inhibition against *C. gloeosporioides* and *P. cinnamomi* was ≥ 20%, using the DNeasy Blood and Tissue kit (Qiagen, The Netherlands). The 16S rRNA gene amplification was performed with primers 27F and 1492R, in a SureCycler 8800 (Agilent Tech., California), with PCR conditions as follows: initial step at 95 °C for 4.5 min; 40 cycles at 95 °C for 1 min, 53 °C for 1 min, and at 72 °C for 2 min, with a final step at 72 °C during 5 min. The amplification of the ITS region was performed with the primers ITS-1 (5′- GTAACAAGGTTTCCGTAGGTG-3′) and ITS-4 (5′-

TTCTTTTCCTCCGCTTATTGATATGC-3′), the PCR cycling parameters were once 95 °C for 3 min, 35 cycles at 95 °C during 45 s, 56.2 °C for 35 s, 72 °C for 45 s, and a final single cycle at 72 °C for 5 min. The DNA amplicons were purified using the Wizard® SV Gel and PCR Clean-Up System (Promega, U.S.A.) and sequenced on a Hitachi 3500 Genetic Analyzer (Applied Biosystems) at ENES-UNAM Morelia. The obtained sequences were deposited in GenBank (accession numbers PV366450 to PV366466 and PV366592 to PV366594). Sequences were manually checked in BioEdit 7.2.5. and aligned using MEGA 7, using the multiple alignment program MUSCLE. The edited sequences and their best matches in GenBank nucleotide database (www.ncbi.nlm.nih.gov) were used to construct the alignment.

### 2.5. Antagonism assays against bee pathogens

The strains of *A. apis* and *P. larvae* were acquired at the National Collection of Microbial Strains and Cell Cultures from the Centro de Investigación y de Estudios Avanzados del Instituto Politécnico Nacional (CINVESTAV).

Antagonism assays against *A. apis* were carried out as described above for the avocado pathogens, using Maltose yeast extract (MYM) plates as medium instead of PDA. To assess the isolateś antifungal activity against *P. larvae*, 200 µL of a standardized suspension (0.5 absorbance at OD600) of *P. larvae* were spread on the surface of Petri dishes with Brain Heart Infusion Agar. Four 5-mm paper discs moistened with 10 µL of standardized suspensions of three nectar microbial isolates and 10 µL of LB medium as a control, were placed at the cardinal points of each plate. After 5 days of incubation at 28°C, the halo of inhibition was measured. Each combination of isolates was tested in triplicate.

### 2.6. Effect of nectar isolates on plant growth and development

*Arabidopsis thaliana* (L., Heynh) ecotype Columbia 0 (Col-0) and reporter line *JAZ1/TIFY10A::GFP:uidA* (Grunewald et al. 2009) were used to evaluate the effect of the selected isolates on plant growth and JA response. *Arabidopsis* seeds were surface-disinfected with 96% ethanol for 5 min and 20% sodium hypochlorite for 7 min. After five washes with sterile distilled water, seeds were stored at 4°C during 48 hours for stratification and grown on agar plates containing 0.2X MS medium (Murashige and Skoog basal salts mixture, Cat. M5524; PhytoTechnology). Plates were placed vertically into a plant-growth chamber (Lumistell ICP-55, Mexico) with a photoperiod of 16 h of light, 8 h of darkness, light intensity of 200 μmol m^2^s^-1^, and temperature of 22°C. Four days after germination (dag), *Arabidopsis* seedlings were transferred to new MS plates, and inoculated with 100 µL (10^8^ CFU, Colony Forming Unit) of the selected isolates.

Control plants were grown without microbial inoculant. Root system architecture (primary root length and lateral root number), biomass accumulation and *JAZ1* expression were analyzed at 7 days after inoculation (dai). The histochemical analysis of transgenic seedlings to evaluate the expression of *JAZ1-GUS* was performed with the β-Glucuronidase Reporter Gene Staining Kit (Sigma-Aldrich), according to the manufactureŕs instructions. Then, plant tissue was clarified according to Malamy and Benfey (1997) protocol, and seedlings were placed on glass slips, covered with coverslips and sealed with commercial nail varnish to analyze J*AZ1* expression with an optical microscope (Zeiss AXIO Zoom V16 with an Axiocam 305 color, Germany).

### 2.7. Metabarcoding and bioinformatic analyses

To characterize the nectar microbial communities, DNA was extracted from 5 µL of each nectar sample using a DNeasy Blood and Tissue Kit (QIAGEN) according to the manufacturer’s instructions. Briefly, from 5 healthy trees we sampled the nectar of 45 female-stage flowers to obtain three composite samples, to construct three libraries for the 16S rDNA and ITS2 metabarcoding; each library was sequenced in triplicate. The preparation of the libraries was performed at the Genomics Lab of the National Laboratory of Ecological and Synthesis (LANASE) following the Illuminás procedure with some modifications. The V4 region of the 16S metabarcode was amplify to characterize the bacterial community, using the primers 515-YF (Parada et al. 2016) and 806R (Appril et al. 2015). For the fungal community, we selected the ITS2 region, using a mixture of four forward primers (Table 1) and the degenerate reverse primer ITS4NGS (Tadersoo et al. 2014) for the amplification. The primers and their sequence are listed in Table 1. The amplicons were indexed with Nextera XT kit v2, and then they were purified with ProNex® Size Selective Purification System following Purification PCR amplicon protocol. Each library was analyzed by capillary electrophoresis with a Qiaxcel system to estimate the final size, 450 bp for the 16S rDNA gene and from 500 to 620 bp for the ITS2 metabarcode, including indexes and adapters. Paired-end sequencing in the format 2 × 250 was performed on a MiSeq instrument (Illumina, USA) at the Sequencing Unit of the Mexican National Institute of Genomic Medicine (INMEGEN). The generated sequences were demultiplexed and the primers removed. Bioinformatics analysis was conducted by QIIME2-2023.7, using DADA2 (Callahan et al. 2016) for trimming and truncating low-quality reads, removing chimeras and generating amplicon sequences variants (ASVs). Taxonomy assignments were performed with the q2-feature-classifer (Bokulich et al. 2018) sklearn with self-trained classifiers. For 16S trained model “SILVA 138 at 99%” database (Robeson et al. 2020) was restricted based on the primers (Werner et al. 2012), and default features were used. The ITS model was trained with full sequences without restriction of UNITE “sh_qiime_release_s_04.04.2024” database (Abarenkov et al. 2024), and k-mer length of [6,6] (Bokulich et al. 2018) and assignments’ confidence limitation of 98% were used. Plastid, mitochondrial and archaeal reads were removed from 16S assignment, and non-assigned reads at phylum level were removed for both datasets. Data derived from bacterial and fungal sequencing were deposited in the Sequence Read Archive of NCBI under accession PRJNA1236352 (https://www.ncbi.nlm.nih.gov/sra/PRJNA1236352).

**Table 1.**
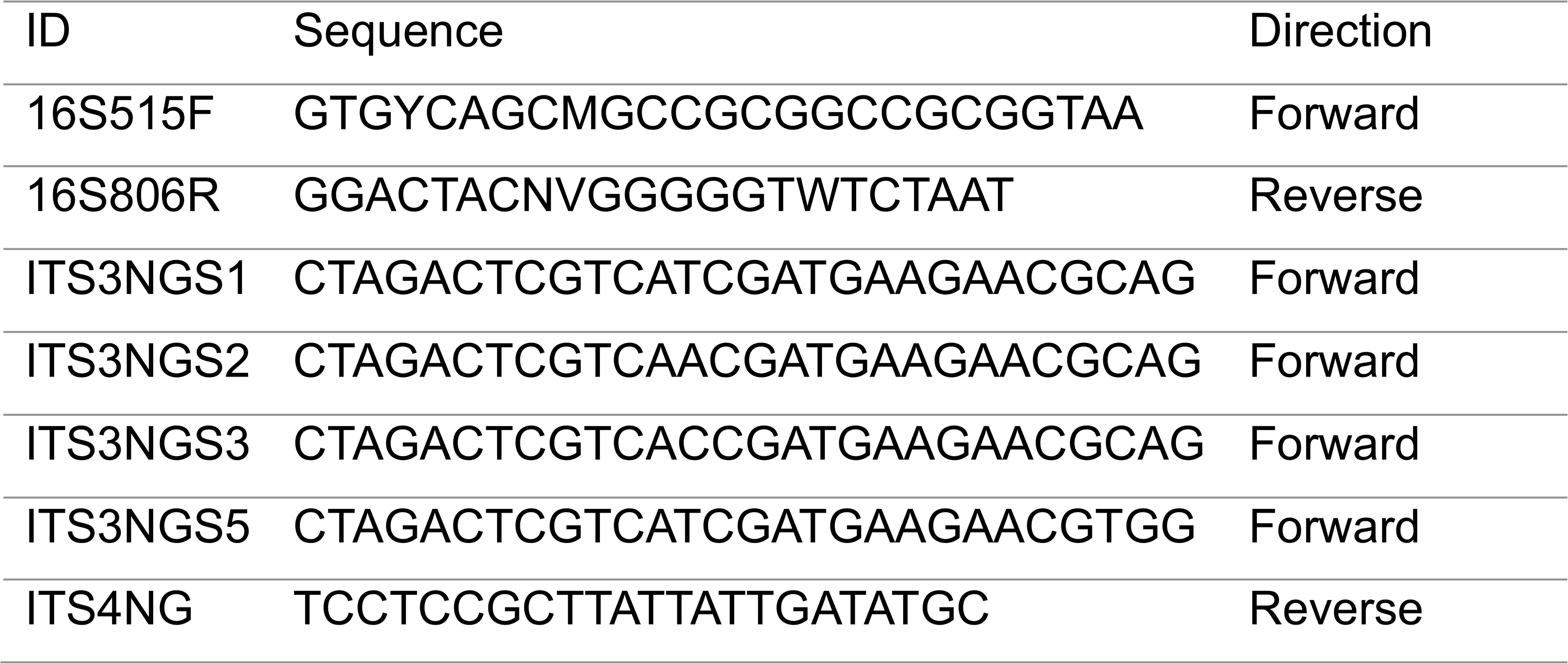
List of primers used to create the 16S and ITS2 gene libraries to characterize the nectar microbial communities.

### 2.8. Statistical analyses

A Shapiro-Wilk test was used to test whether data fitted a normal distribution. A multiple one-way ANOVA with a Tukey’s post-hoc test was conducted to analyze normally distributed data using STATISTICA v.10 software. Treatments were statistically compared to the control using a post-hoc Dunnett test. A t-Student test was performed to determine statistical differences between means of treatments. If the data did not have a normal distribution, a Kruskall-Wallis test was used in R version 3.5.2. For all tests, significance values were (*P*≤0.05).

## 3. Results

### 3.1. Strains isolated from avocado floral nectar differentially inhibit the growth of C. gloeosporioides and P. cinnamomi

A total of 50 flowers from five avocado trees were sampled in the morning right after anthesis, avoiding the arrival of nectar-consuming insects. From the nectar of these 50 flowers, 97 isolates were obtained, which were classified into 43 morphotypes (Table S1); 51 % of them were Gram positive bacteria, 39 % Gram negative bacteria and 9 % yeasts. Representative strains of each morphotype were selected to perform antagonism assays against *C. gloeosporioides* and *P. cinnamomi* by volatile emission or dual culture assays. Two isolates (C-2.5 and H-19.2) showed significant inhibition of the mycelial growth of *C. gloeosporioides* through volatile emission (Fig. 1A), while 17 isolates inhibited radial growth in dual culture assays (Fig. 1A). Isolates I-59.2, B-1.8 and I-70 showed the highest antagonistic activity in dual culture assays, and although their inhibitory effect through the emission of volatiles was not significant, B-1.8 and I-70 caused a lesser mycelium density growth by indirect interaction (Fig. 1B and C). These two isolates were selected to evaluate their effect on hyphae morphology and to perform an avocado fruit rot inhibition assay. Scanning electron microscopy (SEM) micrographs of *C. gloeosporioides* mycelium grown in the vicinity of B-1.8 showed that this strain decreased the thickness of hyphae, causing their collapse and abnormal branching, while I-70 caused hyphal collapse (Fig. 1D). The confrontation of B-1.8 and I-70 against *C. gloeosporioides* by wound and drip in the exocarp of avocado fruits demonstrated that these isolates diminished visibly damage caused by anthracnose (Fig. 1E).

**Figure 1.**
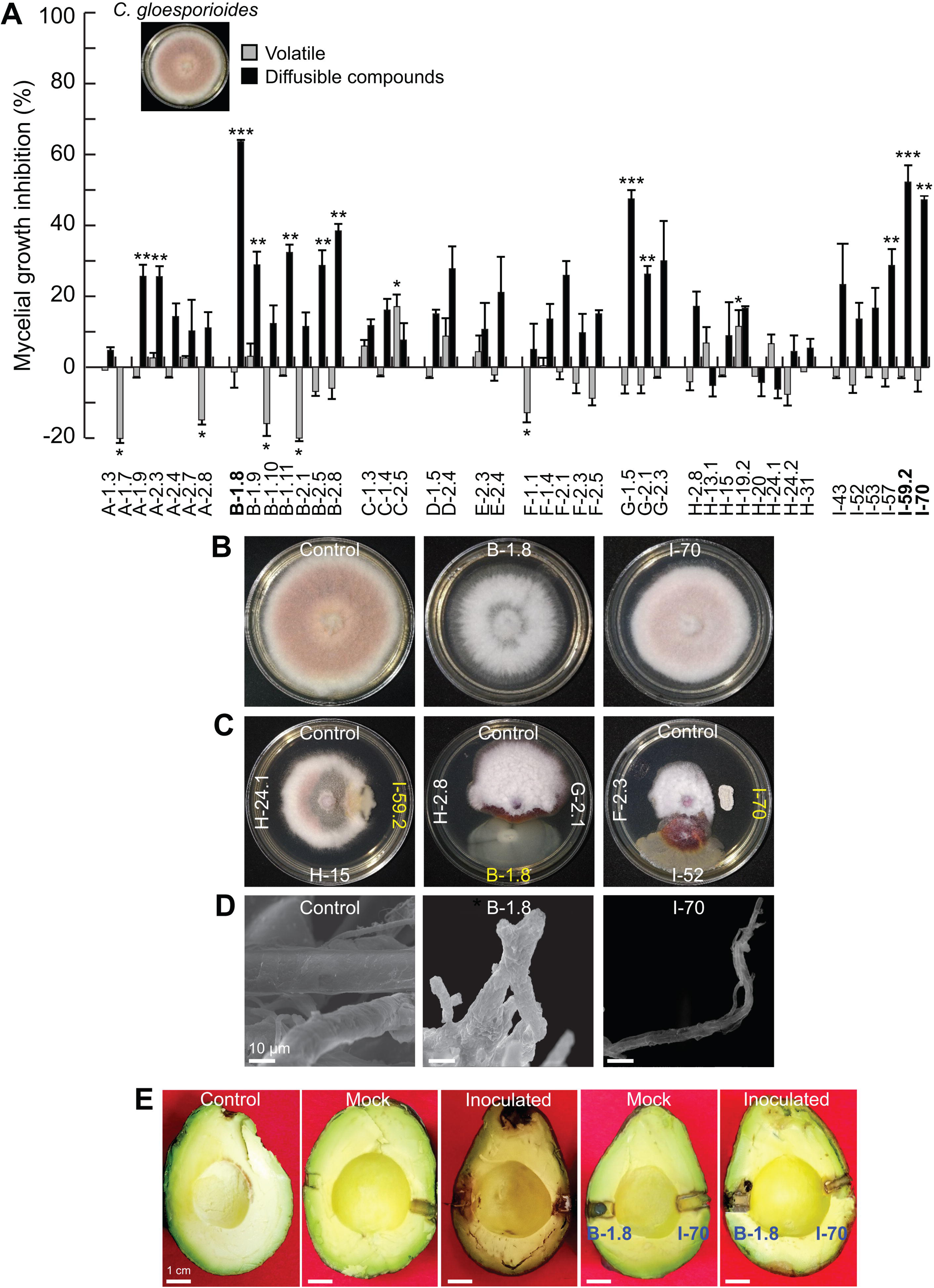
Antagonistic effect of selected microbial isolates from avocado floral nectar against *Colletotrichum gloeosporioides*. Antifungal activity of volatile and diffusible compounds produced by 43 representative morphotypes was evaluated under *in vitro* conditions, and the most active of them were tested in avocado fruit rot inhibition assays. (A) Percentage of inhibition of mycelial growth by emission of volatile and diffusible compounds. (B) Representative images of the mycelial growth inhibition of *C. gloeosporioides*’ plugs exposed to volatiles emitted by the most active microbial isolates. (C) Images of mycelium in dual culture inhibition assays with the most active isolates. (D) Scanning electron microscopy images of the morphological alterations in the hyphae structure induced by the isolates. (E) Representative images of the avocado fruit rot inhibition assays. The values represent the mean ± standard error and asterisks indicate significant difference at *P* ≤ 0.05. All assays were performed in triplicates with similar results.

In the inhibition tests against *P. cinnamomi*, six isolates inhibited the mycelium growth by the emission of volatile compounds and 25 of them showed a percentage of inhibition higher than 20% by direct contact (Fig. 2A). The isolates B-1.8 and I-70 visually reduced the density of the oomycete aerial mycelium by the emission of volatiles and significantly inhibited mycelial growth in dual culture assays (Fig. 2B and C). SEM micrographs showed a drastic reduction in hyphal thickness caused by B-1.8, whilst I.70 induced hyphal thinning and distortion. Both of the isolates caused shriveling of hyphal surface (Fig. 2D).

**Figure 2.**
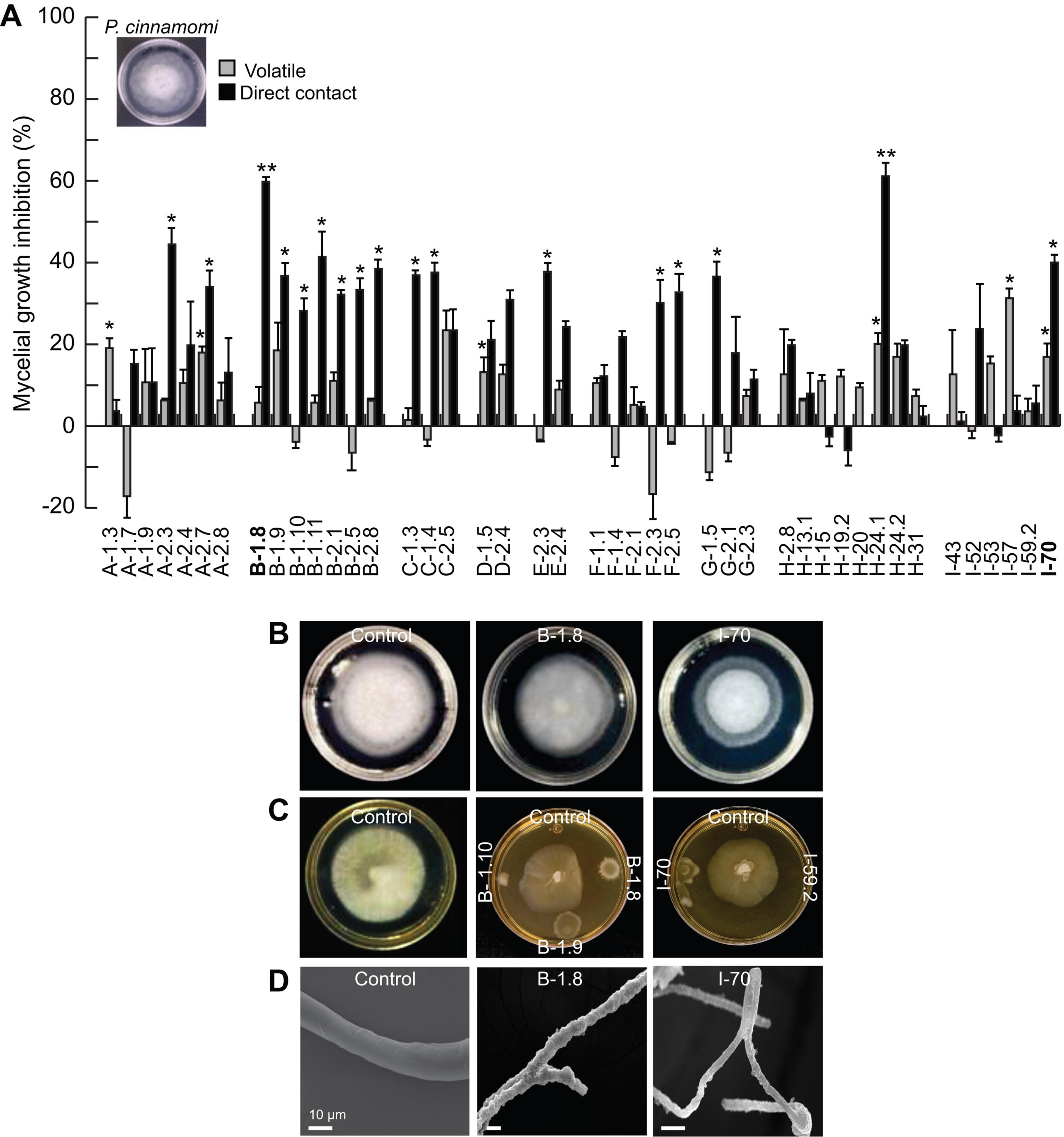
**Inhibition of *Phytophthora cinnamomi* growth by strains isolated from avocado floral nectar. (**A) Inhibition percentage of mycelial radial growth by emission of volatile and diffusible compounds of the microbial strains. (B) Representative images of inhibition assays by volatile emission or (C) in dual culture assays with the most active strains. (D) Scanning electron microscopy images of morphological alterations induced by the isolates with the highest percentage of inhibition. The values shown represent the mean ± standard error, asterisks indicate significant differences at *P* ≤ 0.05. The experiments were repeated three times with similar results.

### 3.2. Molecular identification of avocado nectar strains antagonistic to C. gloeosporioides and P. cinnamomi

The 20 isolates (17 bacteria and 3 yeasts) that were active against the phytopathogens were tentatively identified by 16S rRNA and rDNA-ITS1 - ITS4 gene sequencing. Sequence closest matches based on the BLAST similarity analysis are shown in Table 2. The most represented bacterial genus was *Pseudomonas* with 10 isolates that showed 99-100% identity, being closely related to *P. flavescens* (A-1.9 and A-2.7), *P. aylmerensis* (A-2.3, B-1.8 and F-2.5), *P. protegens* (B-1.9), *P. cerasi* (B-1.10 and B-2.5), and *P. viridiflava* (B-1.1 and G-2.1). Seven of the remaining isolates were closely related to *Paenibacillus favisporus* (B-2.1), *Erwinia aphidicola* (B-2.8), *Curtobacterium ammoniigenes* (C-1.3), *Nocardioides endophyticus* (I-57), *Klebsiella quasipneumoniae* (G-1.5), *Streptomyces eurythermus* (I-70) and *Dietzia timorensis* (I- 59.2). The best match for the ITS1 and ITS4 sequences classified the three yeast isolates into the *Filobasidium* (C-1.4 and G-2.3) and *Aureobasiduim* (D-1.5) genera (Table 2).

**Table 2.**
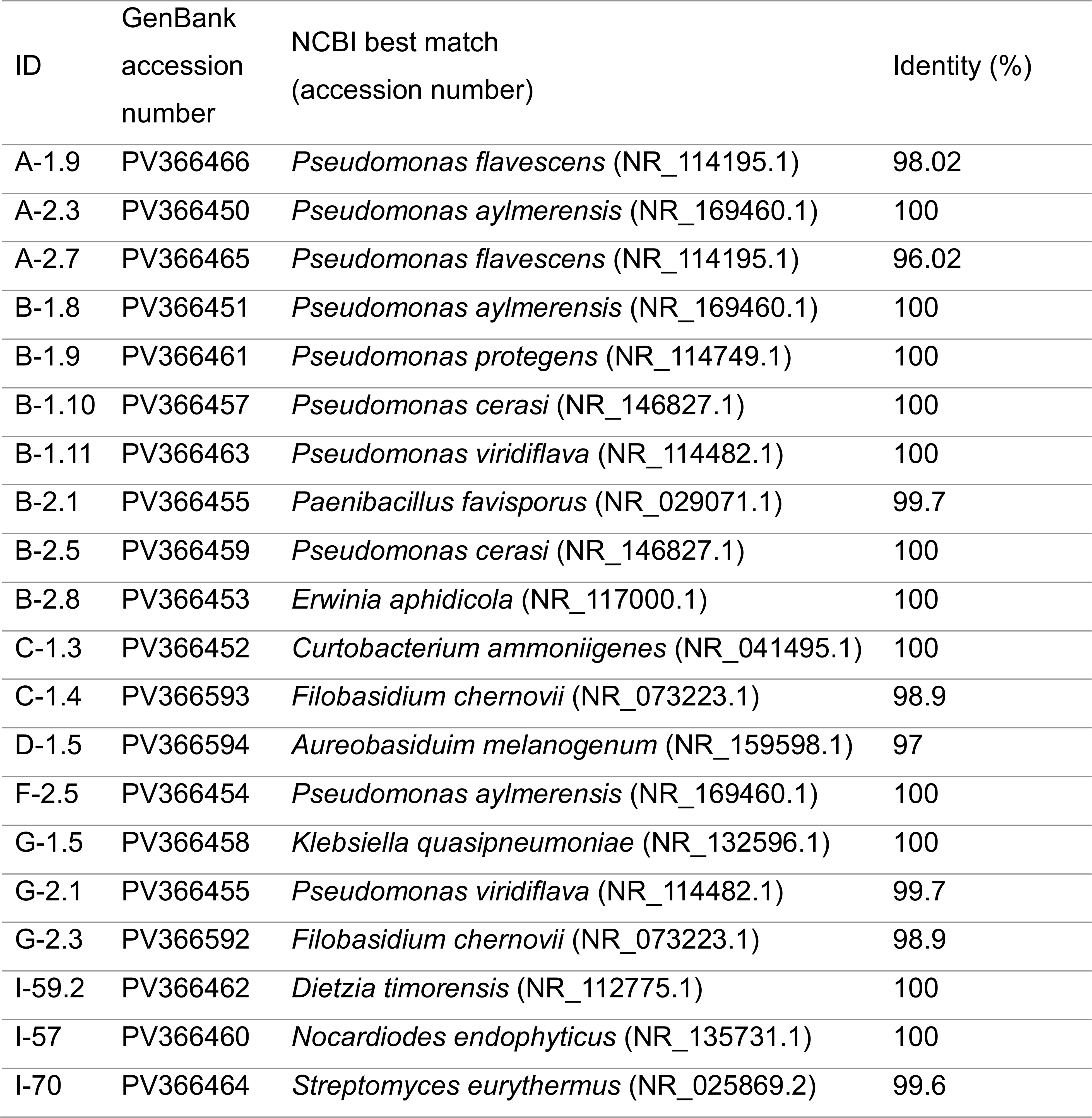
Molecular identification of selected isolates from avocado nectar.

| ID | GenBank<br>accession<br>number | NCBI best match<br>(accession number) | Identity (%) |
| --- | --- | --- | --- |
| A-1.9 | PV366466 | *Pseudomonas flavescens* (NR_114195.1) | 98.02 |
| A-2.3 | PV366450 | *Pseudomonas aylmerensis* (NR_169460.1) | 100 |
| A-2.7 | PV366465 | *Pseudomonas flavescens* (NR_114195.1) | 96.02 |
| B-1.8 | PV366451 | *Pseudomonas aylmerensis* (NR_169460.1) | 100 |
| B-1.9 | PV366461 | *Pseudomonas protegens* (NR_114749.1) | 100 |
| B-1.10 | PV366457 | *Pseudomonas cerasi* (NR_146827.1) | 100 |
| B-1.11 | PV366463 | *Pseudomonas viridiflava* (NR_114482.1) | 100 |
| B-2.1 | PV366455 | *Paenibacillus favisporus* (NR_029071.1) | 99.7 |
| B-2.5 | PV366459 | *Pseudomonas cerasi* (NR_146827.1) | 100 |
| B-2.8 | PV366453 | *Erwinia aphidicola* (NR_117000.1) | 100 |
| C-1.3 | PV366452 | *Curtobacterium ammoniigenes* (NR_041495.1) | 100 |
| C-1.4 | PV366593 | *Filobasidium chernovii* (NR_073223.1) | 98.9 |
| D-1.5 | PV366594 | *Aureobasidium melanogenum* (NR_159598.1) | 97 |
| F-2.5 | PV366454 | *Pseudomonas aylmerensis* (NR_169460.1) | 100 |
| G-1.5 | PV366458 | *Klebsiella quasipneumoniae* (NR_132596.1) | 100 |
| G-2.1 | PV366455 | *Pseudomonas viridiflava* (NR_114482.1) | 99.7 |
| G-2.3 | PV366592 | *Filobasidium chernovii* (NR_073223.1) | 98.9 |
| I-59.2 | PV366462 | *Dietzia timorensis* (NR_112775.1) | 100 |
| I-57 | PV366460 | *Nocardiodes endophyticus* (NR_135731.1) | 100 |
| I-70 | PV366464 | *Streptomyces eurythermus* (NR_025869.2) | 99.6 |

### 3.3. Effects of the phytopathogen-antagonistic strains against Apis mellifera pathogens

The twenty identified isolates were screened for their potential antagonistic activity against the fungus *A. apis* and the bacterium *P. larvae*. Five bacterial isolates and one yeast (C-1.4) showed moderate but significant activity against *A. apis* (*P* ≤ 0.05) by volatile emission; notably, 13 of the tested isolates displayed antifungal activity by direct contact in the dual culture assays (Fig. 3A). From these 13 active strains, *Pseudomonas* spp. B-2.5 and B-1.8, *Streptomyces* sp. I-70, *Paenibacillus* sp. B-2.1 and *Erwinia* sp. B-2.8 inhibited the pathogen mycelial growth by more than 20 % (Fig. 3A-C). In particular, the isolates I-70, B-2.1 and B-1.8, caused hyphal wrinkling and shriveling on *A. apis* (Fig. 3D).

**Figure 3.**
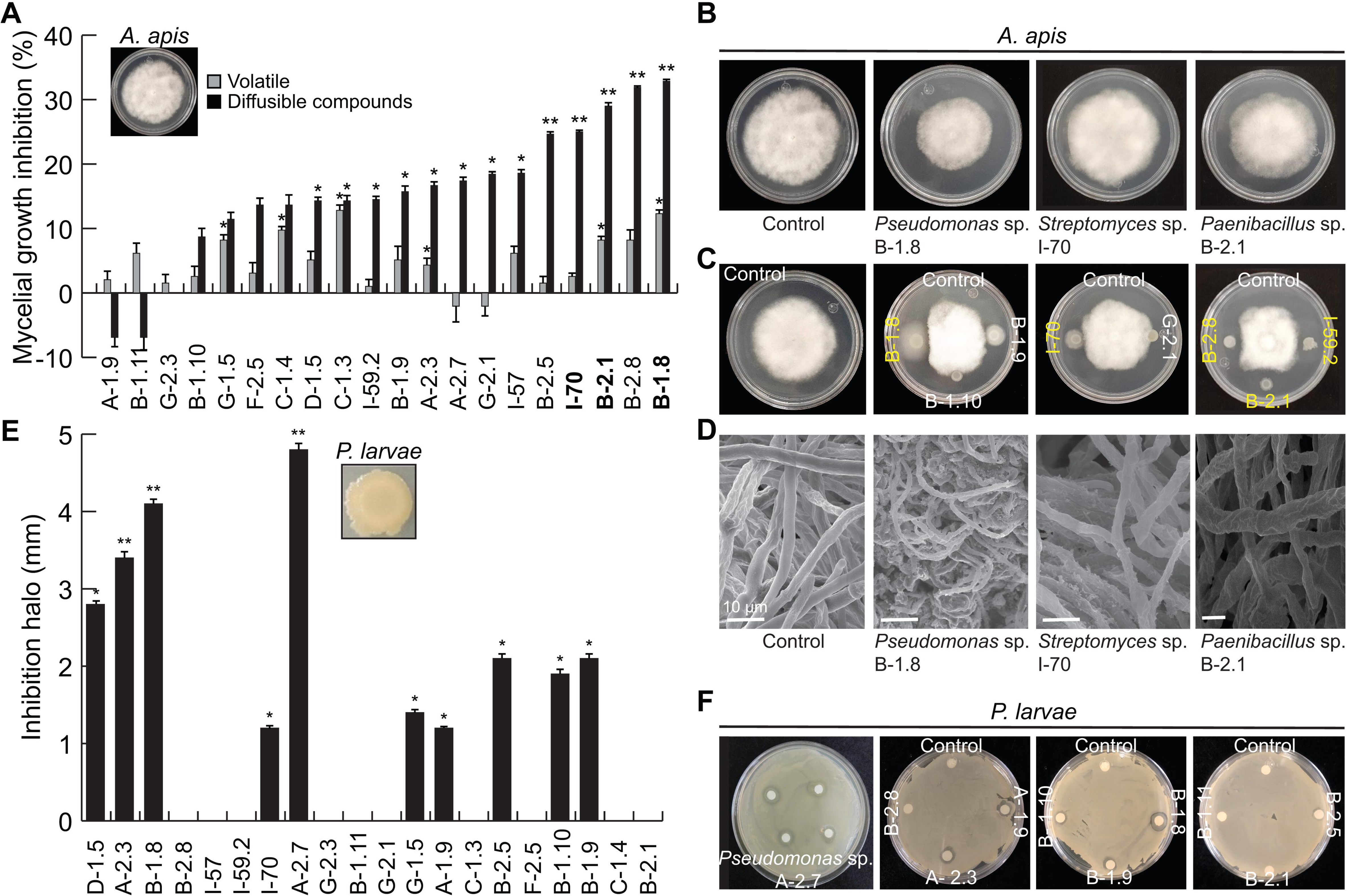
**Effect of the identified isolates on the growth of honey bee pathogens. (**A) Percentage of inhibition of *Ascosphaera apis* mycelial growth by the emission of volatile and diffusible compounds by the selected microbial strains. (B) Representative images of mycelial growth inhibition of *A. apis* exposed to the volatiles released by the most active strains. (C) Dual culture inhibition assays of *A. apis* by the most active strains. (D) Scanning electron microscopy images of the morphological alterations in the hyphal structure of *A. apis* induced by the selected strains. (E) Antibacterial activity of the selected strains co-cultured with *Paenibacillus larvae*. (F) Images of the inhibition assays after five days. The values shown represent the mean ± standard error and the asterisk indicates a significant difference at *P* ≤ 0.05. The assays were performed in triplicate.

Ten of the nectar isolates co-cultured with *P. larvae* generated growth inhibition halos ranging from 1 to 5 mm. *Pseudomonas* spp. A-2.7, B-1.8 and A-2.3 showed the highest inhibiting activity, followed by the yeast *Aureobasidium* sp. D-1.5 (Fig. 3E and F).

### 3.4. Plant growth-promoting properties of avocado nectar isolates

To elucidate whether the 20 selected bioactive strains could have an impact on plant development, we performed *in vitro* interaction assays with *Arabidopsis thaliana*. Outstandingly, the volatiles emitted by all the microbial isolates stimulated growth and development in *A. thaliana* seedlings by increasing the primary root length, promoting the lateral root organogenesis and growth, and increasing biomass accumulation (Fig. 4A-D). When the isolates were co-cultured with the seedlings by direct contact, most of them modified the root architecture; isolates *Pseudomonas* spp. A-1.9, A-2.3 and A-2.7, *Nocardioides* sp. I-57 and *Streptomyces* sp. I-59.2 stimulated root branching or biomass accumulation without any inhibitory effect on primary root length (Fig. 5A-D), while *Pseudomonas* spp. B-1.8 and F-2.5, *Erwinia* sp. B-2.8, *Curtobacterium* sp. C-1.3, and *Klebsiella* sp. G-1.5 decreased the primary root length but stimulated lateral root formation and increased the fresh weight accumulation (Fig. 5A-D). All these data indicate that the avocado nectar microorganisms possess plant growth-promoting activity and that could differentially impact plant signaling pathways.

**Figure 4.**
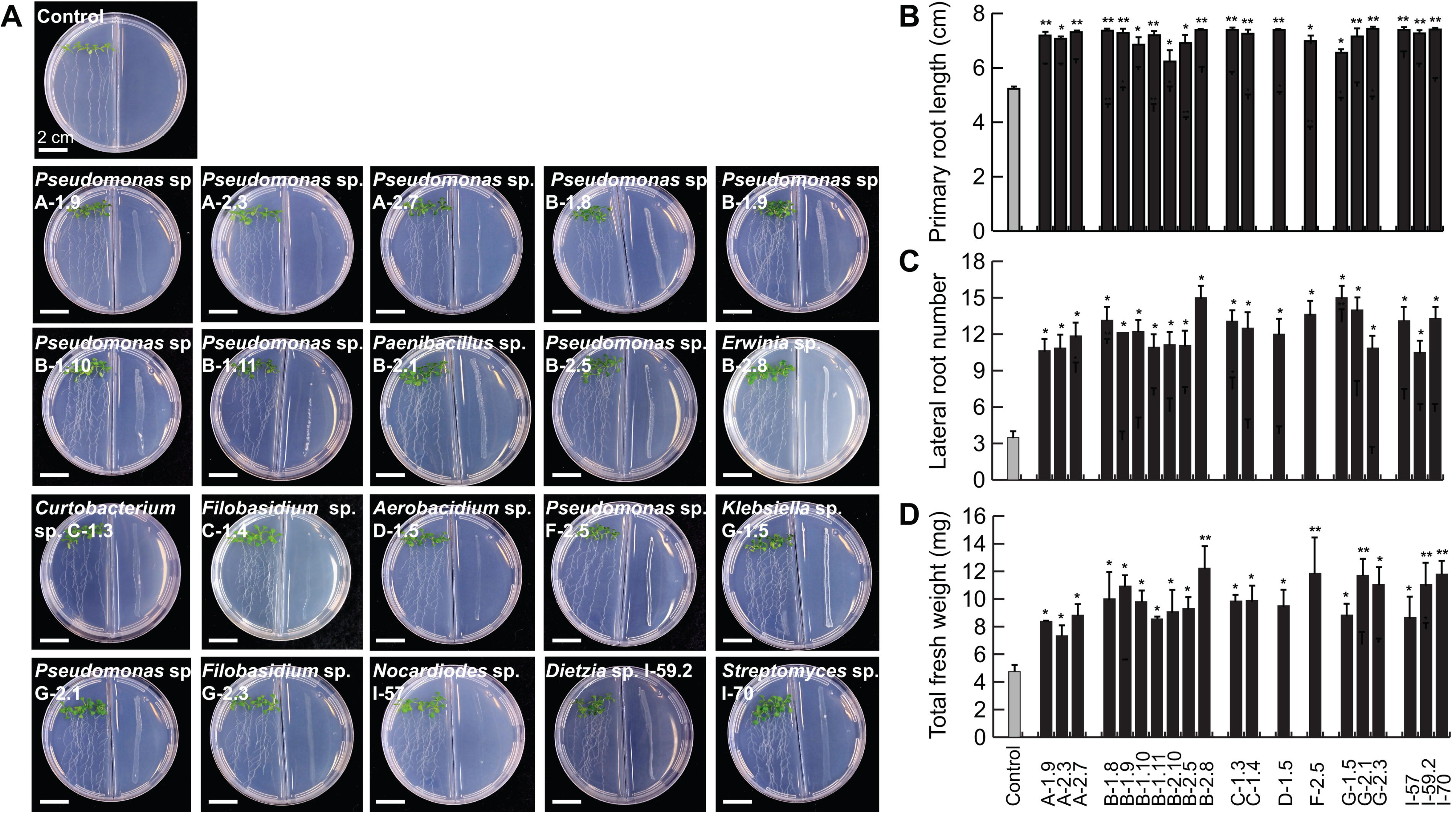
**Effects of volatile compounds produced by the isolates from avocado nectar on the root system architecture and biomass accumulation in *Arabidopsis.*** Four-day-old seedlings were exposed for 7 days to the volatiles released by the selected isolates to determine their effect on plant development. (A) Representative images of *Arabidopsis* seedlings grown in compartmentalized plates in interaction with volatiles produced by the isolates. (B) Quantitative analysis of primary root length, (C) lateral root number and (D) total fresh weight of seedlings grown on the surface of agar MS 0.2× containing plates. Values shown represent the mean ± standard error (n=15 seedlings) and the asterisk indicate a significant difference at *P* ≤ 0.05. The experiments were replicated three times with similar results.

**Figure 5.**
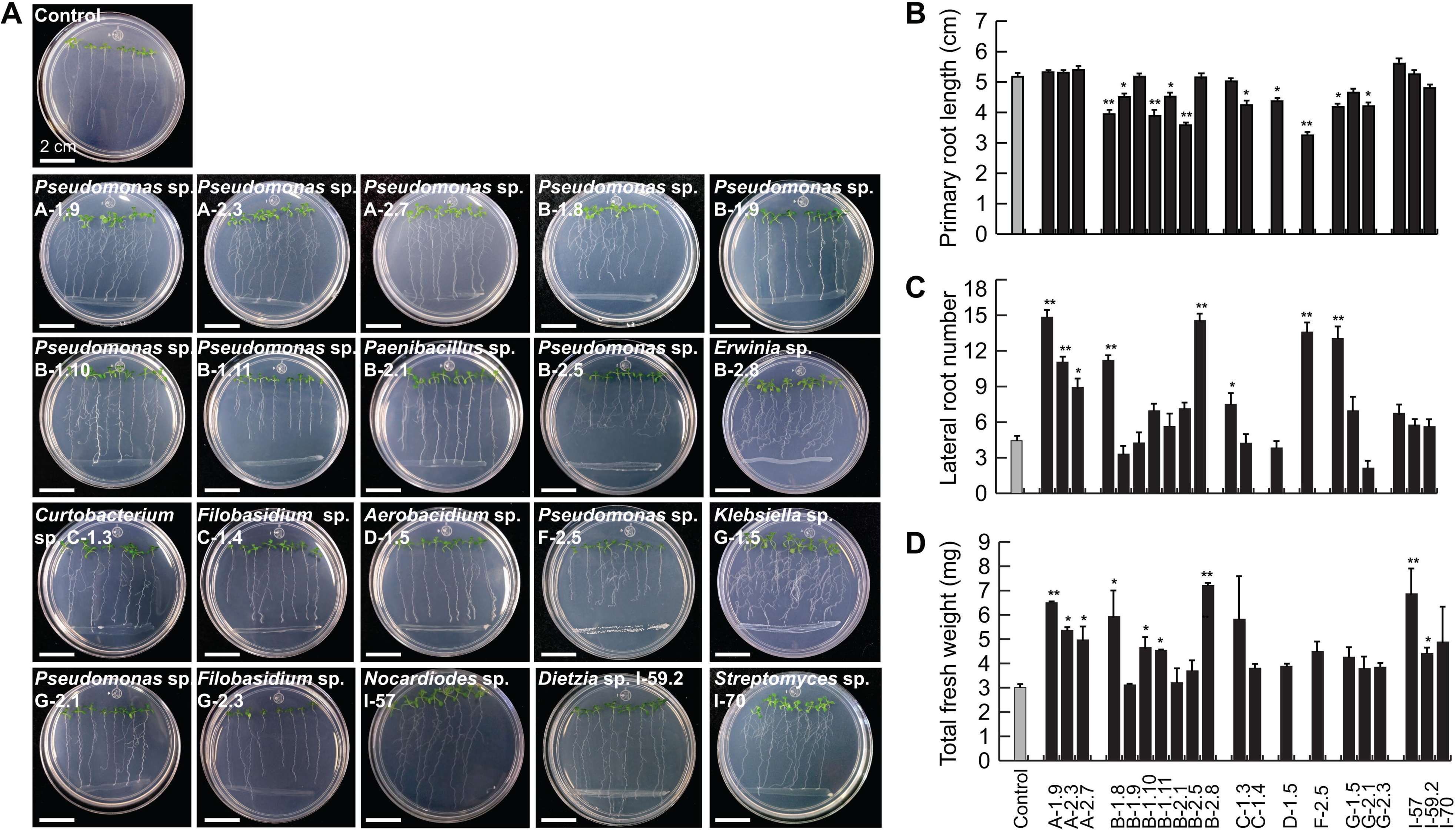
Effects of the co-cultivation of the microbial isolates with *Arabidopsis* seedlings on the root growth and development and biomass accumulation. Four-day-old seedlings were transferred to 0.2× MS medium inoculated with the identified isolates to analyze root architecture and biomass accumulation 7 days after interaction. (A) Representative images of *Arabidopsis* seedlings interacting with the identified isolates. (B) Quantitative analysis of primary root length, (C) lateral root number and (D) total fresh weight. Values shown represent the mean ± standard error (n=24 seedlings) and asterisks indicate a statistically significant differences compared to the control (*P* ≤ 0.05). The experiment was repeated in triplicate with similar results.

### 3.5. Induction of defense responses in A. thaliana by the plant growth-promoting isolates

Jasmonic acid (JA) is one of the main phytohormones orchestrating the plant defense response against biotic stressors with developmental programs (Raya-González et al. 2012; Carvalhais et al. 2017). Sensing of plant growth-promoting (PGP) microorganisms may result in triggering the JA-mediated induced systemic resistance (ISR) that defends plants from herbivores and necrotrophic pathogens such as *C. gloeosporioides* and *P. cinnamomi*. Therefore, we analyzed the expression of the JA-responsive *Jasmonate-zim domain protein 1* (*JAZ1*) gene as an indicator of pathway activation in *JAZ1/TIFY10A::GFP:uidA A. thaliana* transgenic seedlings, in direct contact with the 20 strains selected for their activity against necrotrophic phytopathogens and PGP properties. Histochemical GUS activity assays indicated that *Pseudomonas* spp.

A-1.9, A-2.3, B-1.8, B-1.11, F-2.5 and G-2.1, *Paenibacillus* sp. B-2.1, and *Sreptomyces* I-70 induced an exacerbated gene expression in shoot tissues and in the apical region of primary roots, from the xylem to the external cell layers, which coincides with developmental shifts such as root hair differentiation, shortness of meristematic zone and thickening of roots (Fig. 6). The bacterial isolates *Pseudomonas* spp. A-2.3, A-2.7 and B-2.5 and *Curtobacterium* sp. C-1.3, and the yeasts *Filobasidium* spp. C-1.4 and G- 2.3, and *Aerobasidium* sp. D-1.5 induced the expression of the JA-responsive gene only in the shoot system (Fig. 6), probably indicating the activation of a ISR response. The isolates *Erwinia* sp. B-2.8, *Klebsiella* sp. G-1.5, *Dietzia* sp. I-57, *Nocardioides* sp. I-59.2 and *Streptomyces* I-70 induced the systemic activation of the JA-responsive pathway with a moderated root expression and shoot induced signal of the gene marker (Fig. 6), which could be differentially related with a PGP or with a pathogenic activity.

**Figure 6.**
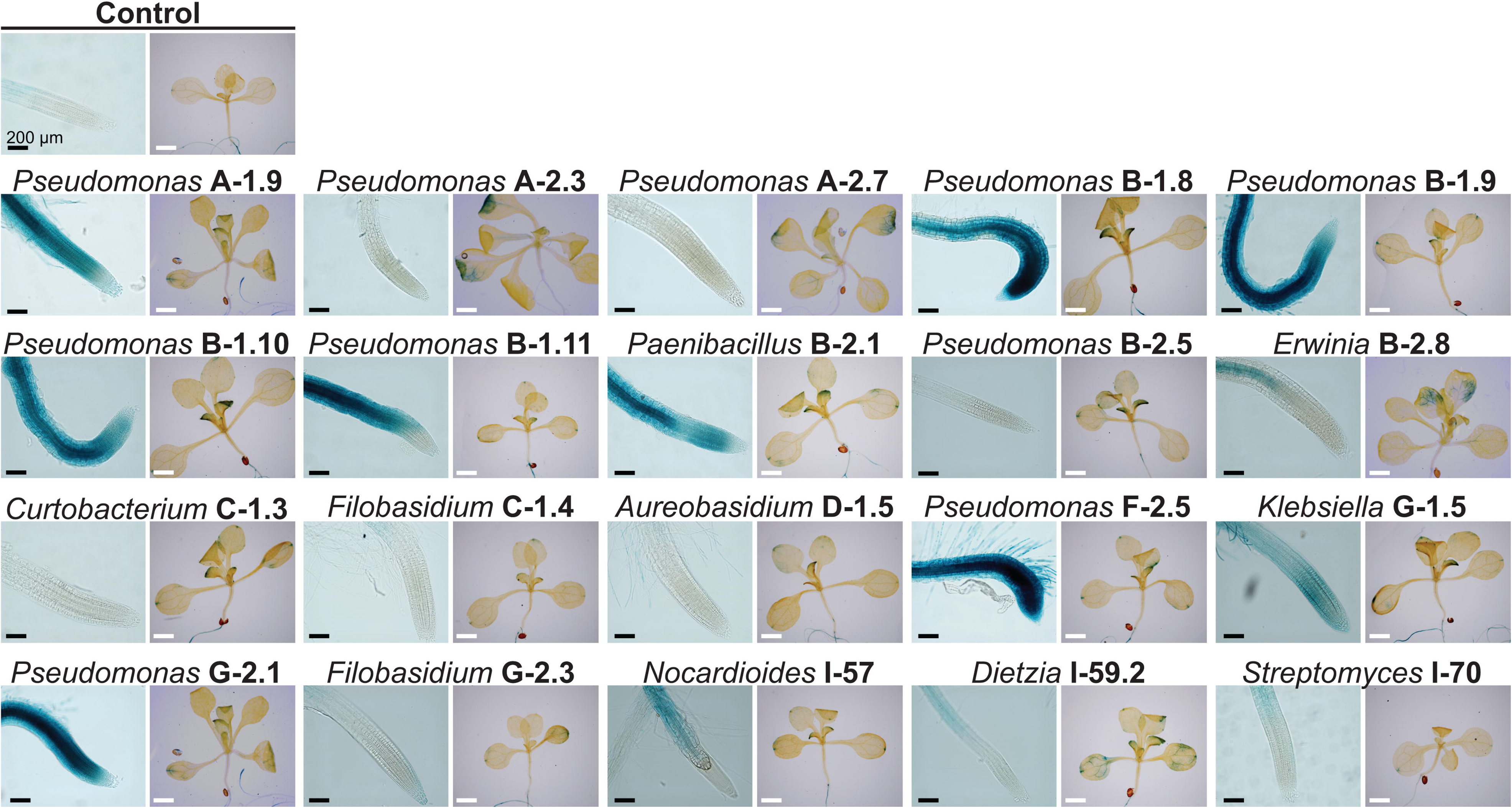
Effects of selected isolates on defense-related jasmonic acid gene expression in *Arabidopsis*. Four-day-old transgenic seedlings expressing the jasmonic acid inducible *JAZ1/TIFY10A::GFP:uidA* gene marker were transferred to 0.2× MS medium inoculated with the different isolates to analyze the activation of JA signaling pathway 7 days after the interaction (n = 24 seedlings).

### 3.6. Microbial community composition in the avocado floral nectar

We employed a metabarcoding approach to explore the complexity of the nectar microbial community and to assess how representative of its diversity could be the cultivable fraction previously described. A total of 634 bacterial amplicon sequence variants (ASVs) and 249 fungal ASVs, with a minimal size of 234 bp after filtering with DADA2 were recorded. The rarefaction analysis revealed that the abundance of ASVs reached a plateau, indicating that the saturation of sequencing depth for the samples was reached, except for the triplicate 3A in the 16S sequencing. The dominant bacterial order was Pseudomonadales, represented by the *Acinetobacter* and *Pseudomonas* genera, reaching a relative abundance from 40 to 94%, followed by *Massilia* and *Polaromonas* associated to Burkholderiales, and by *Sphingomonas* (Fig. 7A and Fig.

**Figure 7.**
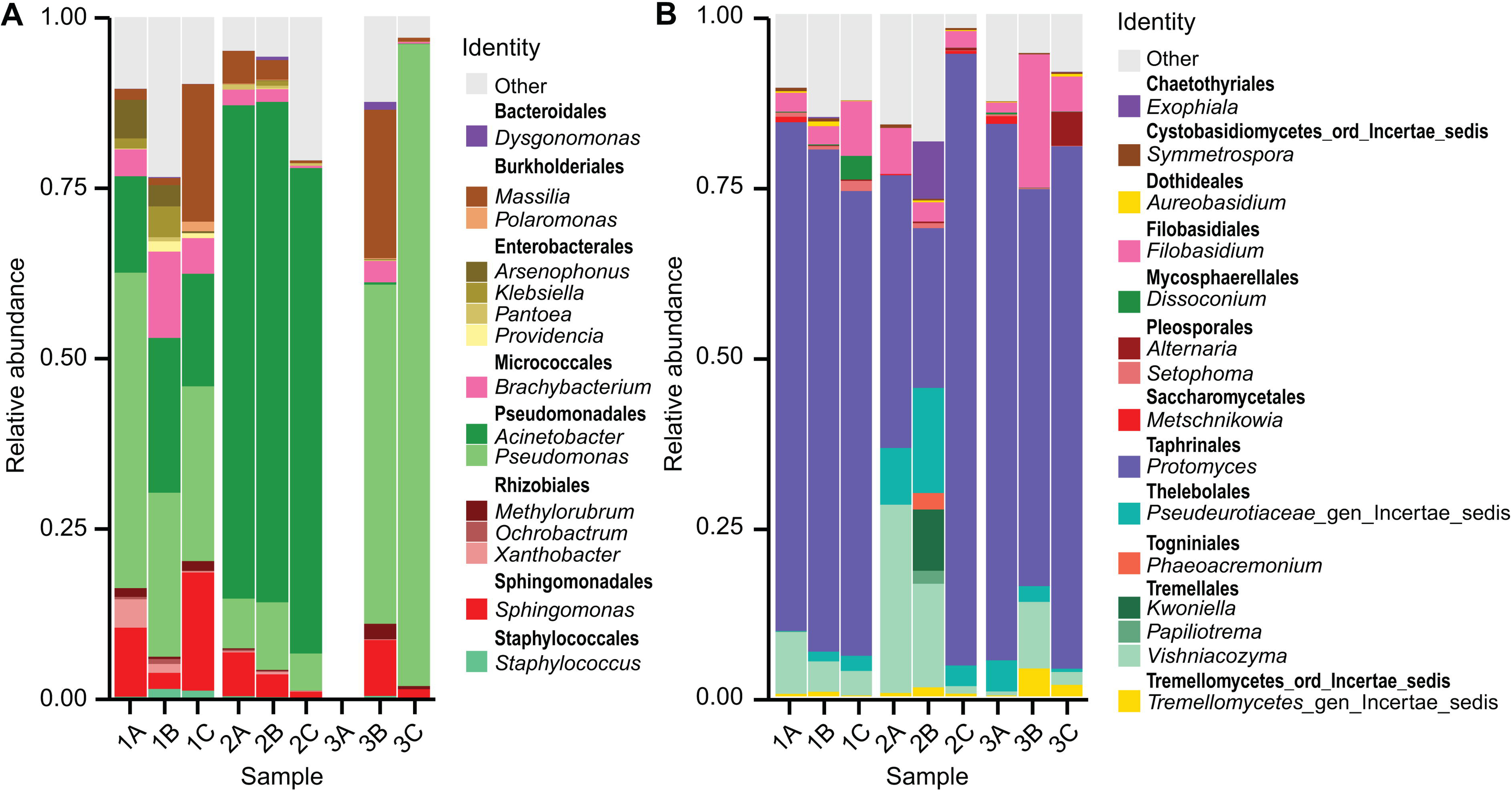
Structure of the microbial community associated with the avocado floral nectar. Relative abundance of (A) bacterial and (B) fungal genera obtained by Metabarcoding sequencing. A total of 634 bacterial ASVs with a 99% confidence limitation and 243 fungal ASVs with a 98% confidence limitation were assigned.

S1). The most abundant fungal ASVs were *Protomyces* (23 – 89%), the phyllosphere-inhabiting *Vishniacozyma* (0.6 – 27%) and *Filobasidium* (1.5 – 19.5%) (Fig. 7B and Fig. S2). The low diversity and large dominance recorded for the bacterial and fungal communities from nectar samples evidence the high selectivity that avocado nectar exerts over the microbial community.

## 4. Discussion

In angiosperms, floral structures constitute the first interphase in plant-pollinator interactions, where nectar represents the main nutritional reward (Vannette, 2020). Floral nectar harbors a specialized microbiota that could impact both plant and pollinator health (Vannette et al., 2013; Martin et al., 2022; Álvarez-Pérez et al., 2024). However, beneficial traits of the nectar microbiota to antagonize plant and pollinator pathogens and promote plant fitness at early developmental stages have been scarcely explored.

Here, we reported for the first time the composition of the nectar-inhabiting bacterial and fungal communities, and some functions of their culturable fraction, in a crop with agronomical, ecological and evolutionary importance, the avocado *Persea americana* Mill.

Our data indicated that 20 out of 43 tested isolates were active against *C. gloeosporioides*, *P. cinnamomi* and entomopathogens in dual culture, whilst their emitted volatiles promoted plant development. This reinforces the theory that the few microbial taxa present in nectar play important roles for plant fitness (Good et al., 2014) and demonstrates a high potential of nectar for bioprospecting beneficial microorganisms for agriculture. Most tested isolates displayed a larger pathogen inhibition through direct contact than through volatile emission, which suggests that the inhibition of mycelial growth observed in dual culture assays could be due to the production of diffusible compounds and cell wall degrading enzymes. *Pseudomonas* spp., *Paenibacillus* spp. *Klebsiella* spp., *Aureobasidium* spp. produce phenazines, exophilins, lyamocins and lytic enzymes such as proteases and chitinases that damage the fungal cell wall (Merín et al. 2014; Raza et al. 2015; Di Francesco et al. 2017; Jisha et al. 2018; Hsu et al. 2021; Zhang et al. 2022; Iqbal et al. 2023). This was reinforced with the microscopy images of the mycelium of the different pathogens; the analysis of the metabolomic and volatile profiles produced by our antagonistic microbial isolates should be further studied.

Regarding the effect of nectar microorganisms on plant fitness, microbial volatile and direct contact interactions with our model plant showed that volatile emission by all the evaluated strains promoted root branching and biomass accumulation, while the accumulation of some diffusible microbial metabolites led to activation of the JA defense pathway. Volatile and diffusible compounds from *Pseudomonas* spp*. Klebsiella* spp*., Dietzia* spp.*, Streptomyces* spp., and *Aureobasidium* spp. have been explored for their role in promoting plant development. Benzyl alcohol, 2-phenylethyl alcohol, and 3- (methyl- thio)-1-propanol, trimethylamine, 3-hydroxy-2-butanone (acetoin), 2,3- butanediol, 2-pentylfuranare are some examples of VOCs produced by these genera that promote nutrient availability and root development (Camarena-Pozos et al. 2019; Jones et al. 2019; Ravelo-Ortega et al. 2023) and should be further studied for their effects on avocado development. In this context, the activation of JA signaling by nectar microbial isolates is of particular interest, since it could, in addition to coordinating the defense against necrotrophic fungi, drive nectar secretion and the plant response against herbivores and their mechanical damage (Heil et al. 2012; Millán-Cañongo et al. 2014; Wang et al. 2021; López-García et al. 2023). These JA -inducing properties are specially striking in terms of ecological interactions, since some pathogenic bacteria can move into the vascular system of the host plants through the floral nectaries, i.e. *Erwinia tracheiphila*, the causative agent of wilt disease in cucumber (Sasu et al. 2010). A previous report described how the strawberry rhizospheric strain *Streptomyces globisporus* SP6C4 is able to translocate endophytically from roots to flowers and back to the roots, and to antagonize the gray mold pathogen *Botrytis cinerea* whilst reducing honeybee mortality caused by *P. larvae* and *Serratia marcescens* (Kim et al. 2019).

These results support the idea that the rhizosphere, the anthosphere and their associated microbiota impact the plant ecosphere, including pollinatorś health. Future work should strive to evaluate the capacity of avocado nectar microbes to colonize the plant endosphere and stimulate developmental and defense related pathways.

It is important to note that avocado nectar communities may also possess other functional attributes that are not explored here. According to several reports, some isolates belonging to the genera *Dietzia* and *Nocardioides* could protect plants against salt stress, heavy metals and other air pollutants (Bharti et al. 2016; Ma et al. 2023; Shahid et al. 2024). Some species related to our identified isolates have traditionally been reported as phytopathogens, such as *C. flaccumfaciens* yields (López-Ramírez et al. 2022; Tokmakova et al. 2024) and nectar specialists *Erwinia* (Pozo et al. 2011; Jacquemyn et al. 2013; Vannette, 2020). However, our data indicated that volatile and diffusible compounds of *Erwinia* sp. B-2.8 and *Curtobacterium* sp. C-1.3 promoted root system development in *A. thaliana* seedlings.

Molecular identification of the selected isolates largely coincided with the genera that reached the highest relative abundances in the metabarcoding analysis. *Pseudomonas* was one of the most dominant genera with a relative abundance of 32.8%, and 50% of the bioactive isolates belonged to this genus. In a similar way, *Erwinia*, *Klebsiella* and *Filobasidium* were among the taxa with the highest relative abundance in the nectar microbial community, and were also retrieved in the culturable fraction. However, there were exceptions with *Paenibacillus sp.* B-2.1, *Curtobacterium sp.* C-1.3, *Dietzia* I-57*, Nocardioides* I-59.2, and *Streptomyces* I-70, which were not detected among the most abundant taxa in avocado nectar, and *Acinetobacter* that was the bacterial genus with the highest relative abundance, although none of the 20 isolates selected for molecular identification corresponded to it. This could be because *Acinetobacter* spp. from avocado nectar do not present antagonistic activity against *C. gloeosporioides* and *P. cinnamomi*, which was our main selection criterion. However, since antagonistic activity of *Acinetobacter* spp. strains has been reported against several *Colletotrichum* and *Phytophthora* species, this warrants future efforts to retrieve *Acinetobacter* isolates and investigate their potential bioactivity (Syed-Ab-Rahman et al. 2018; Patel et al. 2019; Matos et al. 2024).

Ubiquitous microbes reach nectar via wind, rain, and animal transport. However, only a few taxa are able to survive UV radiation, the presence of antimicrobial compounds, low nitrogen availability, and osmotic stress that characterize nectar as a highly selective ecological niche (Pozo et al. 2011). A low microbial diversity in nectar community composition suggests that dominant taxa could be particularly important for plant health (Good et al., 2014; Jacquemyn et al., 2021). In hexose-rich nectars predominate specialist yeasts of genus *Metschnikowia*, followed by the generalist *Aureobasidium* and *Cryptococcus*, while the most common bacteria belong to the Enterobacteriaceae family and to the genera *Acinetobacter* and *Bacillus* (Pozo et al. 2011; Vannette, 2020; Mueller et al. 2023; Álvarez-Pérez et al. 2024). Here, the high relative abundance of *Acinetobacter*, *Pseudomonas* and *Vishniacozyma* in avocado nectar was consistent with that reported in other nectars, but *Bacillus*, *Metschnikowia* and *Cryptococcus* were not included among the most abundant taxa in the analyzed samples, while this was the first report of *Protomyces* as the dominant taxon in nectar. These differences in communities are probably due to the chemical composition of avocado nectar and the species that pollinate its flowers (de Vega et al. 2009). Perseitol content in avocado nectar could be inhibiting the germination of *Bacillus* spores (Rizk et al. 1989), while being a nectar rich in sucrose with low fructose and glucose content is not optimal for the development of an alcohol-fermenting yeast incapable of hydrolyzing sucrose, such as *Metschnikowia* (Vicente et al. 2020; Torrellas et al. 2023). The case of *Protomyces*, a little-researched phytopathogen in floral nectar with the ability to assimilate sucrose and which causes hyperplasia and formation of galls in the veins of leaves, petioles, peduncles, stems and fruits (Čadež et al., 2021; Leharwan et al., 2021), is interesting, since there are no reports of this genus as a causal agent of diseases in avocado (González-Santarosa et al. 2014), suggesting that avocado nectar could act as a reservoir for this phytopathogen. Finally, even the most abundant genera are likely to benefit from each other, since not all have the ability to use sucrose as a carbon source. *Acinetobacter* spp. and *Filobasidium* spp. are non-saccharolytic, the bacteria have n-alkanes, as their sole carbon sources and the yeast degrade higher fatty acid esters and terpenoids to promote the aroma and flavor in fruits (Wang et al. 2023; Wu et al. 2023). They likely thrive on avocado nectar at the expense of n-alkanes, VOCs and polyphenols synthesized to mediate interactions with insects and provide protection against UV radiation, bacteria, and fungi as occurs in other plant species (Bartwal et al. 2013). On other hand, *Aureobasidium* spp. and *Pseudomonas* spp. can assimilate a wide range of compounds, but the later prefers organic acids and amino acids as a carbon source, rather than sugars (Rojo, 2010; Chi et al. 2022), therefore they could benefit from the tricarboxylic acids produced by other microorganisms that inhabit the nectar (Peng Wang et al. 2022; Anderson et al., 1985).

Collectively, our results show that avocado nectar harbors a competitive specialized microbiota with bioactivity either as phytostimulant, through the emission of volatile compounds, or as contributors to the plant health, by antagonizing pathogens and activating the JA defense pathway. Our findings suggest that the nectar microbiota could directly impact the plant fitness and contribute to its pollinatorś health. Future work will focus on exploring the ability of the nectar-living microbes in avocado to colonize the plant host and to disperse in the environment.

## 5. Concluding Remarks

The composition and some ecological functional traits based on culture-independent and -dependent on the floral nectar microbiota here described in *Persea americana*, an ancient angiosperm with a high agroalimentary value, represent a fundamental precedent to examine the mechanisms of early co-evolutive adaptations of flowering plants and the assembly of their associated microbiomes. Collectively, our findings highlight the selectivity of avocado floral nectar over its inhabiting microorganisms, their potential beneficial effects in the avocado-honeybee pollination system as antagonistic agents against plant and bee pathogens, and their ability to stablish belowground interactions with plants.

## Acknowledgements

This research was funded by the Mexican Council of Humanities, Science and Technology (CONAHCYT-SECIHTI), grant number CB-2014, 242999 and by the CONAHCYT-PRONACES-PPE grant (project “PERSEA”, 322772). C.M.L.-G. thanks Dirección General de Asuntos del Personal Académico (DGAPA, UNAM) for her posdoc fellowship grant and A.M.-B. (CVU 50123) the fellowship grant (854328) received from CONAHCYT- SECIHTI during his sabbatical leave. We would like to extend our sincere gratitude to Mr. Alfonso Barragan, owner of the orchard “Rancho La Joya” for kindly allowing us to sampling during the flowering season. Our special acknowledgments to Hernando Rodríguez Correa for his technical assistance and logistics support, to Silvana Martén Rodríguez for her advising and facilitating materials for the nectar sampling, to Valentina García Aguilar for her technical support and to Javier Raya González and Arturo Guevara García for the kind donation of *Arabidopsis* seeds. We are also thankful to Valerie Martin and Aurélie Deveau for the enriching and critical discussions during the writing of this work.

## Notes

### Competing Interest Statement

The authors have declared no competing interest.

