## Supplementary figures and images for "Floral nectar microbiota of *Persea americana* inhibits pathogens and improves plant fitness"

### Figure S1

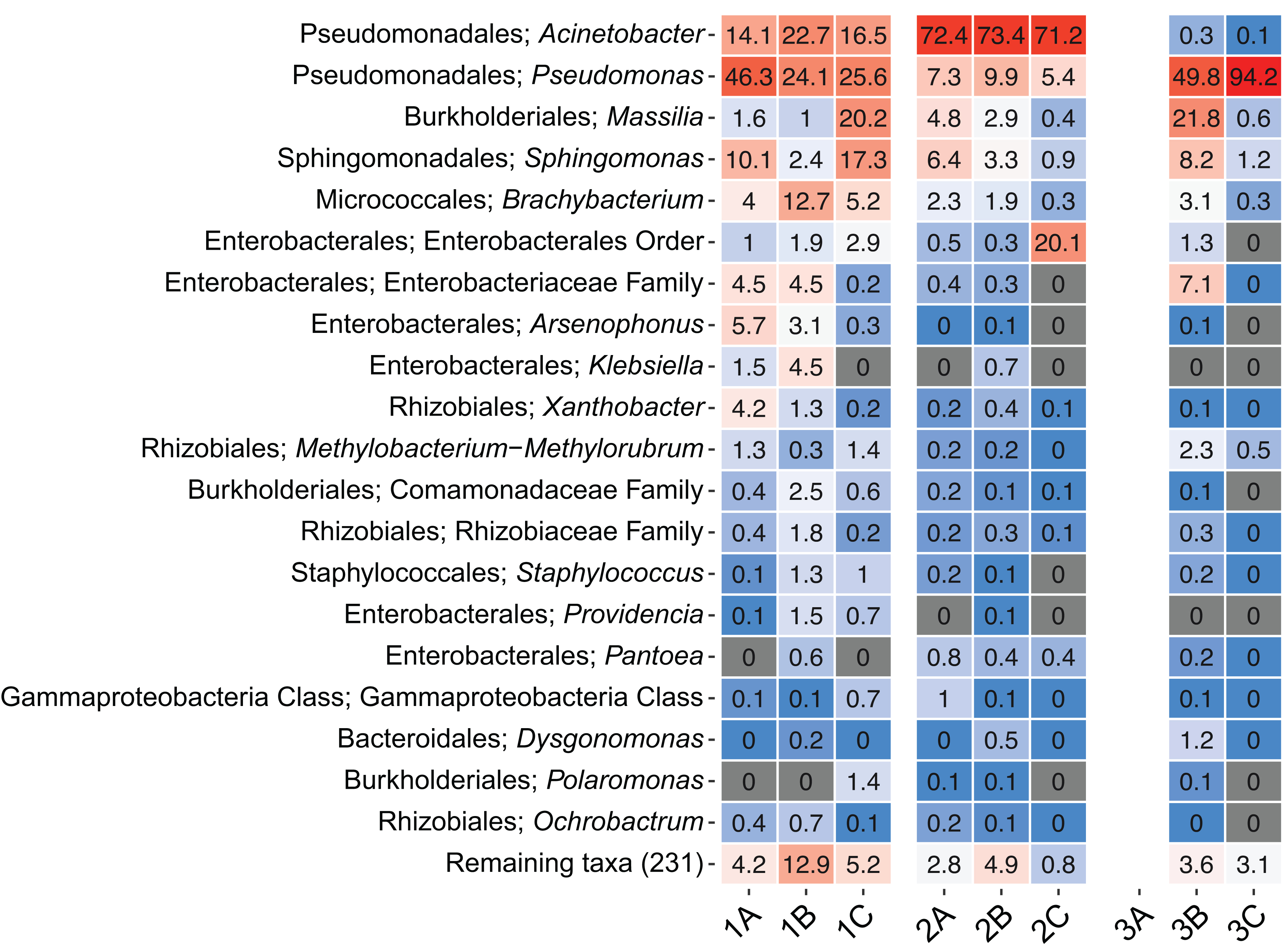

### Figure S2

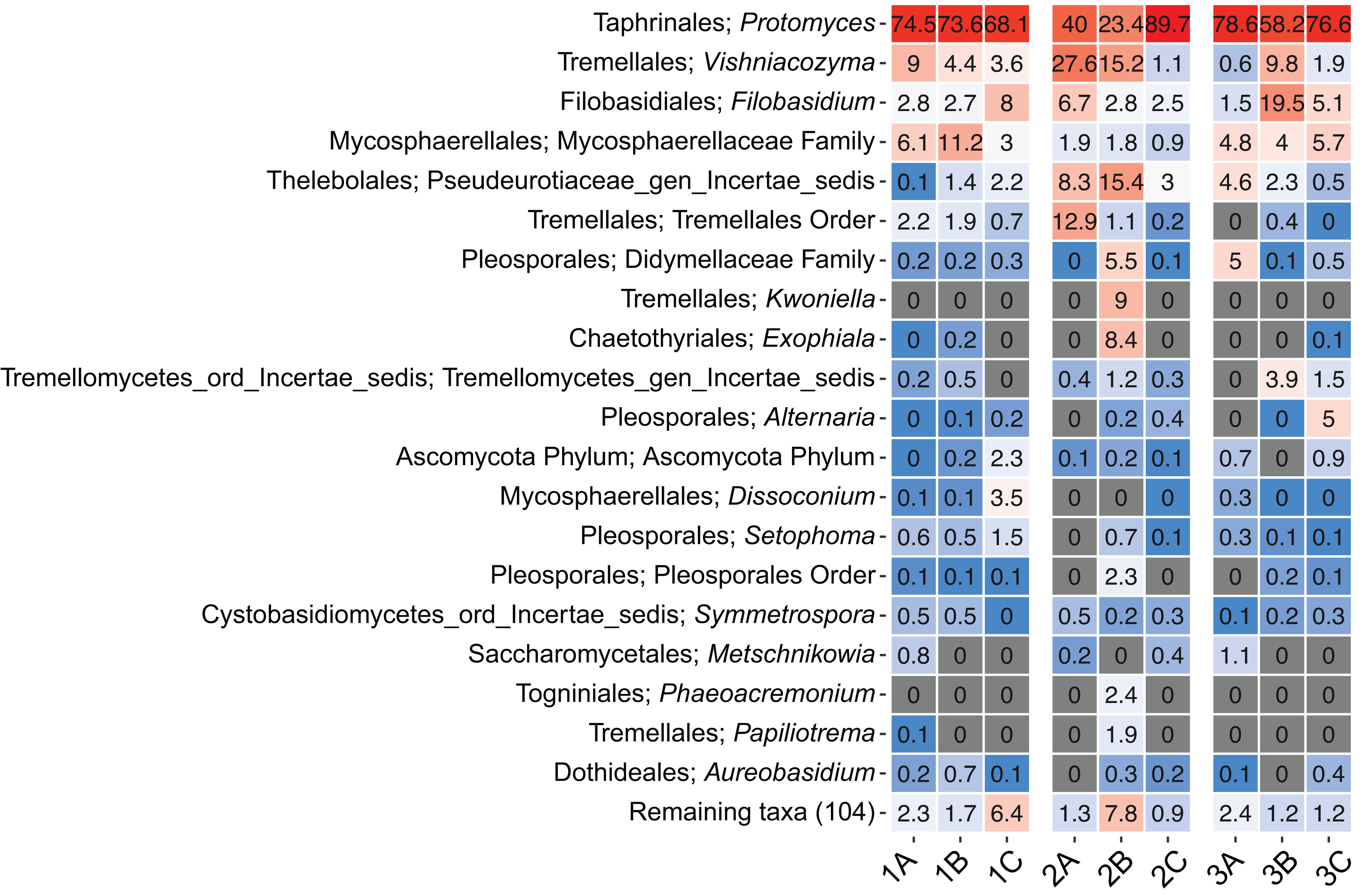
